# Real-time AI integration for MR to detect artifacts and guide pulse sequence adaptations

**DOI:** 10.64898/2026.05.04.722724

**Authors:** Aaron T. Gudmundson, Zahra Shams, Abdelrahman Gad, Shuyuan Wang, Dunja Simicic, Saipavitra Murali-Manohar, Gizeaddis L. Simegn, Ipek Özdemir, Christopher W. Davies-Jenkins, Yulu Song, Vivek Yedavalli, Georg Oeltzschner, Omer Burak Demirel, Jeremias Sulam, Michael Schär, Sandeep Ganji, Richard A. E. Edden

**Author notes:** Corresponding Author: Aaron T. Gudmundson, Ph.D. The Malone Center for Engineering in Healthcare 600 N. Wolfe St, Park 367, Johns Hopkins University, Baltimore, MD, USA – 21287-0005.

## Abstract

This work presents a first-of-its-kind artificial intelligence (AI-)integrated MR pulse sequence that detects out-of-voxel (OOV) artifacts in real-time (within-TR) and responds prospectively by updating the crusher gradient scheme.

Per Excitation Real-time Execution & Guided Responses with Integrated Neural-network Evaluation (PEREGRINE), allows for deployment of deep learning models and pulse sequence updates. In this study, PEREGRINE operated a time-domain (TD) and frequency-domain (FD) convolutional autoencoder that detect OOV artifacts. Scans without (AI-off) and with (AI-on) updates were collected from the medial prefrontal cortex of healthy volunteers using a MEGA-edited MRS experiment. The degree of OOV contamination (OOV Score) was quantified per transient based upon the prevalence of OOV signals in the TD and FD data. OOV Scores above a user-defined threshold triggered an update of the crusher gradient scheme, iterating through 48 permutations (6 axis transpositions × 8 polarity flips).

Within each 2-second TR, PEREGRINE successfully provided single-transient OOV Scores and updated gradients accordingly. No difference was observed between the OOV Scores from the full (“Full” condition) AI-on and AI-off sessions due to the AI-on scan cycling over better and worse gradient permutations relative to the AI-off scan. However, the AI-on scan had significantly lower OOV Scores than the AI-off scan when selecting the transients where PEREGRINE persisted (“Dwell” condition) on a given gradient permutation. Ultimately, Fit Quality Number (FQN) from linear combination modeling improved significantly for the AI-on compared to the AI-off scan.

PEREGRINE enabled an adaptive and AI-integrated sequence allowing for real-time evaluation and response to OOV artifacts, identifying gradient modifications that produced less OOV contamination.

**Graphical Abstract:** *Graphical Abstract Summary:* PEREGRINE, Per Excitation Real-time Execution & Guided Responses with Integrated Neural-network Evaluation, allows for pulse sequence updates using on-scanner deployed neural networks. To demonstrate the utility, PEREGRINE was used to operate two neural networks (time/frequency domain), trained to detect out-of-voxel artifacts, during a MEGA-edited MRS experiment in the prefrontal cortex. When artifacts were detected, PEREGRINE updated the crusher gradients within the same TR. Real-time, adaptive, and AI-driven MR will provide a long-awaited solution for combatting artifacts and poor data quality.

*Graphical Abstract Figure:* 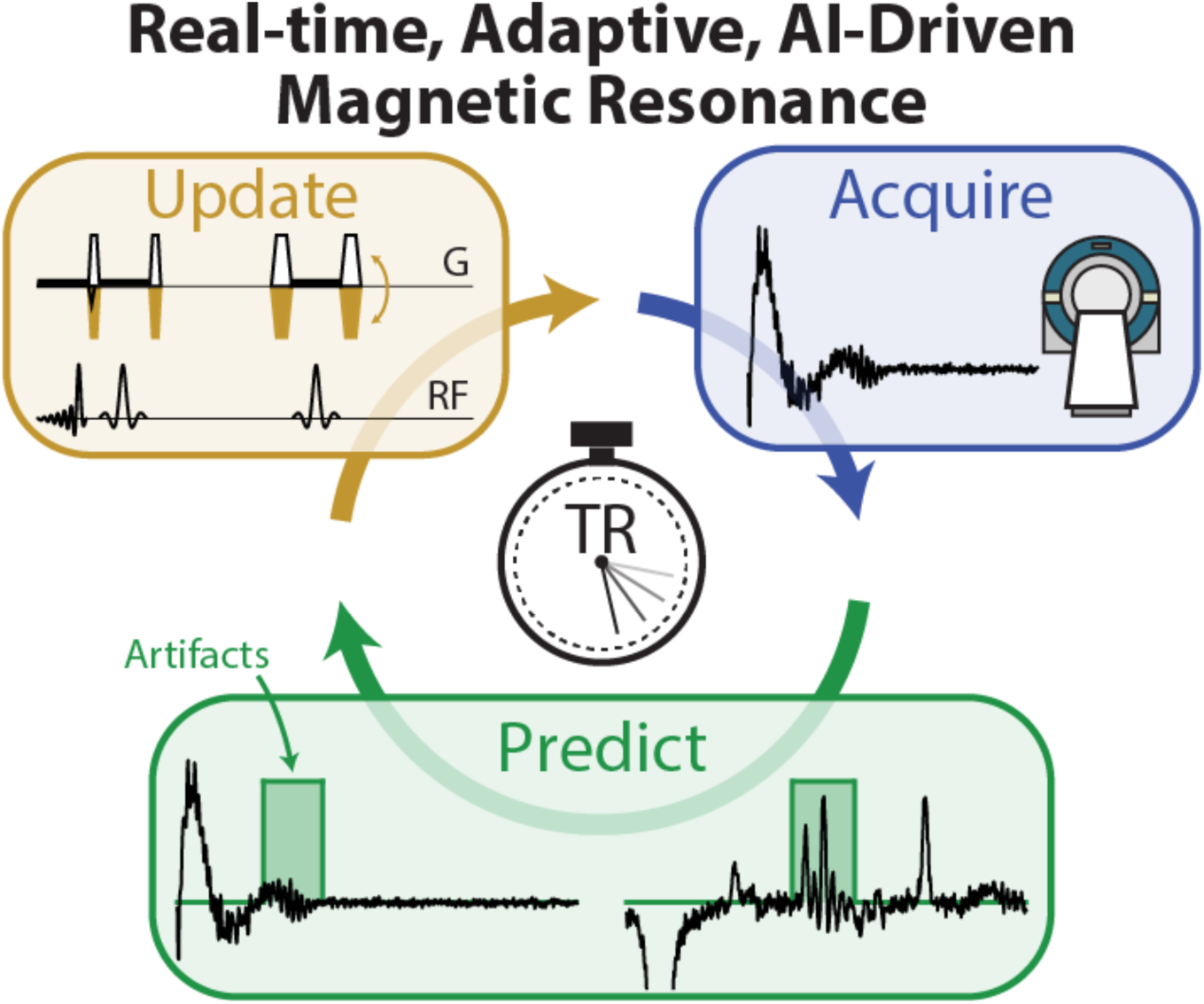

## 2. Introduction

Magnetic resonance spectroscopy (MRS) is a non-invasive technique that can measure neurochemicals in vivo within the brain. Commonly measured metabolites include neurotransmitters (e.g., GABA and glutamate), antioxidants (e.g., ascorbate and glutathione), and energy metabolites (e.g., creatine and lactate)^1,2^. Metabolite signals have low amplitude relative to thermal noise, so MRS experiments often last several minutes, collecting more than100 transients. Metabolite signals are also small, relative to potential nuisance signals, including those from tissue water and sub-cutaneous lipids. Therefore, MRS experiments require careful preparation (e.g., field homogeneity, RF calibration, etc.) and are susceptible to artifacts associated with subject motion and scanner instability. Data analysis for MRS usually occurs after the subject has left the scanner, on a separate workstation, with the artifacts usually only recognized when they cannot easily be addressed.

Experiments that are long, unstable, and artifact-prone can benefit from acquisition methods that update acquisition parameters prospectively during the experiment itself. Motion correction is the most common corrective update, using interleaved navigator scans^3,4^ or external optical tracking^5–7^ to determine head position, and update voxel location (in the scanner frame) to maintain the target location (in the subject frame). More advanced corrective methods can update shim fields, typically using dual-echo navigators to map the B0 field and stabilize F0 and field homogeneity within the voxel^8–13^. More recently, it has been proposed that MRS experiments might be updated not just in response to auxiliary navigators, but *to the MRS data themselves*^14^. An adaptive multi-echo relaxation approach was demonstrated to reduce experimental time without sacrificing measurement sensitivity – the echo times sampled were adjusted in response to the data being collected using a classical Bayesian model^14^.

Artificial intelligence (AI) through deep neural networks has shown immense promise for signal processing^15–23^, artifact detection^24–27^, and quantification^28–30^ in MRS. Neural networks are ideal for real-time adaptive MRS, because they are effective in solving ill-defined problems with ambiguous solutions^31^, and the inference of a trained neural network is fast^32–34^ allowing for TR-to-TR updates. The incorporation of AI in an adaptive scanning framework has the potential to ameliorate common problems that lead to poor quality MRS data. To demonstrate the potential of deep learning for corrective updates, a use case should be considered which includes a poorly defined problem space, sequence-level updates, and the potential to benefit from real-time, within-TR updates.

Out-of-voxel (OOV) artifacts are a persistent problem in MRS^35,36^ and can severely degrade the quality of MRS data. These artifacts are currently only identified during data analysis, and their solution remains uncertain. OOV artifacts primarily arise from unsuppressed water signals outside the voxel of interest, that are unintentionally refocused as echoes by local field gradients^35,37,38^. Such artifacts are often problematic when measuring lower-concentration metabolites, and using sequences with more pulses (e.g., edited sequences)^39^. OOV artifacts are most catastrophic in brain regions with high B0 inhomogeneity, such as near air-tissue interfaces (e.g., sinuses or ear canals), and for smaller voxels, as the out-of-voxel frequency dispersion increases due to the reduced shim volume, resulting in less efficient frequency-selective suppression and larger regions of unsuppressed water signal.^35,37,38,40^. OOVs overlapping with a signal of interest may bias spectral modeling resulting in uncertainty in measurement, and/or exclusion of the poor-quality data.

Four main methods are used in MRS experiments to suppress unwanted signals: gradient selection; phase cycling; frequency-selective suppression; and spatially selective suppression. Ideally, the gradient scheme should dephase signals that originate from outside of the voxel^41^, but hardware constraints (e.g., gradient amplitudes, higher order shim gradients) and experimental design (e.g., TE constraints) limit their effectiveness. Optimized gradient schemes^42–44^ offer some benefit, but because gradient selection harnesses spatially dependent phase to suppress unwanted signals, any signal thus suppressed might be refocused by a local gradient of the correct direction and sufficient amplitude^38,39^. Phase cycling selects for the desired coherence pathway by averaging across transients^45,46^, but imperfect pulses, motion, field instabilities and long cycle lengths lead to incomplete cancellation of unwanted signals. Frequency-selective water suppression^47,48^ does not remove OOV signals because they arise from water in regions where B0 is shifted (by 10s to 100s of Hz)^38,40^. Spatial suppression by ‘saturation bands’ can help, but the location of an OOV artifact is unknown a priori, so this is not a reliable strategy^49,50^. Recent work has started to characterize the spatial and pathway origin of OOV signals^39,51^, which will, in turn, improve the application of all suppression methods.

The framework from Starck et al., 2009 considers OOV signals as being encoded to a particular starting point in k-space by the sequence gradients, with the local field gradient performing the role of the read-out gradient in an imaging experiment to form a gradient echo^38^. The direction of evolution in k-space during the read-out is determined by the local field gradient direction, and the starting point, by the relative directions, amplitudes and polarities of the sequence gradients. As suggested within this framework, any individual OOV signal can be suppressed by flipping the polarity of all sequence gradients, since its starting point in k-space would be inverted and its direction of evolution in k-space during read-out would then be away from the k-space origin so that no gradient echo occurs. Unfortunately, OOV signals that were previously suppressed may be refocused by this new gradient scheme, and the OOV problem has a ‘whack-a-mole’ character. When this process of OOV detection and experiment adjustment occurs once-per-experiment, it is not possible to make much progress with a subject- and voxel-specific issue. Deep learning has demonstrated a strong ability to identify OOV artifacts^24,26^, eliminating the need for local expertise and automating the process of flagging poor-quality data. These neural networks have performed well on single transients, reducing the length of time needed to make an evaluation and demonstrating their potential for online processing at the scanner.

Here, we introduce PEREGRINE (Per Excitation Realtime Execution & Guided Responses with Integrated Neural-network Evaluation) to make pulse sequence adaptation based upon neural networks that are deployed directly on the scanner. In this study, PEREGRINE was used to operate neural networks to individual transients, testing for the presence of OOV artifacts. If OOV artifacts were found, PEREGRINE changed the gradient scheme in real time to suppress them.

## 3. Experimental

The following sections describe in detail the training, implementation, and deployment of the AI-integrated pulse sequence to address OOV artifacts. Upon acquisition of each transient of the sequence, data are pre-processed and analyzed to return an ‘OOV Score’ (described in detail in section 3.2.). If that OOV Score exceeded a preset threshold, the data quality is deemed insufficient, and the pulse sequence gradient scheme is updated for the subsequent TR, following the flow diagram shown in Figure 1.

**Figure 1.**
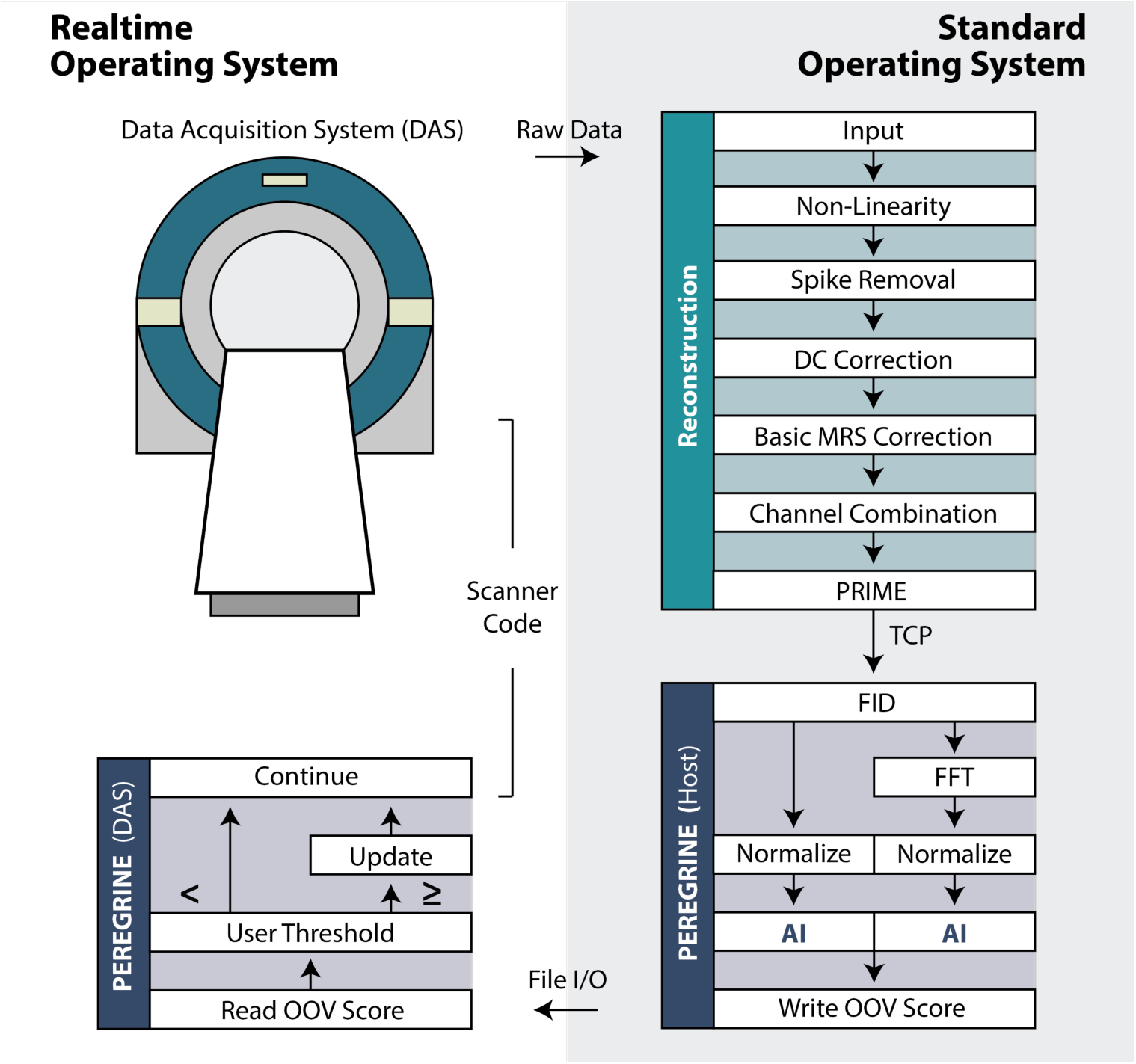
Diagram of the Realtime Workflow. Data collected on the spectrometer, or DAS, running the VxWorks RTOS is passed to the Reconstructor on the Windows SOS. Following Channel Combination, data are intercepted and sent to the PEREGRINE workflow with a custom implementation of the Philips Reconstruction Injector Modulator and Emitter, or PRIME, reconstruction node^57^. Neural networks are run on the SOS, and the result is read back to the RTOS to update the pulse sequence. This workflow is repeated once every two-second TR, with the sequence elements consisting of 1,826 ms (722 ms water suppression + 80 ms TE + 1,024 ms acquisition time) and an additional 174 ms delay during which PEREGRINE executes.

### 3.1. Neural Network

In line with previous work^24^, the AI methodology relied entirely upon synthetic data to train two neural networks for complex-valued time-domain (TD) and frequency-domain (FD) input. Specifically, the AGNOSTIC synthetic dataset, which consists of metabolite, residual water, macromolecules, and noise components each corresponding to one of 18 field strengths and 15 echo times, was used. As described previously^24,52^, metabolite components were generated from a linear combination of basis functions derived from highly-sampled time-domain density matrix simulations with metabolite amplitudes informed by Reference 52. Additional OOV artifacts modeled with 5 parameters (center index, width, frequency, phase, amplitude; according to Equation 1 of Reference 24). OOV artifacts were restricted to between 0.0 ppm and 4.1 ppm to obstruct the target metabolites. 191,400 examples were used with 180,000 for training, 1,800 for validation, and 9,600 for testing. Ground-truth binary masks for each example were determined as all points above 5% of the maximum amplitude of the Gaussian OOV signal envelope.

The two neural networks used the same fully convolutional autoencoder architecture as the previous work^24^, but were developed using PyTorch to take advantage of the LibTorch C++ API^53^. The neural networks were both trained as before^24^, this time, receiving either a time-domain FID or frequency-domain spectrum and producing an OOV detection vector of the same size 1 × 1 × 2048 × 2 (batch, channels, datapoints, real/imaginary). The OOV detection vector is a binary mask with 0 indicating no OOV artifact and 1 indicating the presence of OOV artifact for a given point. The Dice coefficient^54–56^ was used to measure the overlap between the OOV detection vector and the ground-truth binary mask for the loss function during training and to evaluate performance during testing.

### 3.2. Per-Excitation Realtime Execution & Guided Responses with Integrated Neural-network Evaluation PEREGRINE

The PEREGRINE framework for the AI-integrated pulse sequence was implemented on a 3 Tesla MR scanner (Achieva dStream, Philips, Best, The Netherlands), and included custom software for data reconstruction, neural network deployment, scanner GUI modification, and pulse sequence integration. This software included a combination of code for the WindRiver VxWorks Real-Time Operating System (RTOS) that is used to operate the precisely timed spectrometer and the Microsoft Windows Standard Operating System (SOS) that is used for data reconstruction and the user interface for scan preparation.

PEREGRINE deploys the trained neural networks and serves as the continuous link between FIDs from the reconstruction environment and updates to the pulse sequence on the spectrometer with both host-side and spectrometer-side components (See Figure 1). PEREGRINE uses LibTorch (PyTorch API) to deploy the networks in C++ with minimal latency. PEREGRINE is initialized at the beginning of the scan session, and the networks are loaded into CPU memory (CPU, to avoid unnecessary CPU-to-GPU data transfer time as the single-transient data would not benefit from GPU parallelization). Throughout the scan session, PEREGRINE provides log statements using a command prompt to inform the user of successful delivery of raw data, live neural network results, and give confirmation the results are available for the spectrometer. The full process is described below.

After acquisition of each individual transient, data are transferred from the embedded RTOS of the spectrometer, or Data Acquisition System (DAS), to the reconstruction environment on the SOS. Upon complete transfer, data are moved through a series of reconstruction nodes to generate a time-domain FID. The reconstruction pipeline is designed for internal libraries, and requires a separate application to incorporate third-party AI dependencies. Therefore, data are intercepted within this pipeline and transferred using a custom implementation of the Philips Reconstruction Injector Modulator and Emitter (PRIME) reconstruction node^57^ which uses Transmission Control Protocol (TCP) to stream data and a header with necessary metadata (e.g., number of data points, field strength, spectral width, etc.).

Upon arrival to PEREGRINE, a copy of the FID is Fourier transformed (Cooley–Tukey algorithm^58^) to maintain both time- and frequency-domain representations of the transient. Next, each domain is normalized, reshaped, and converted to a tensor (torch::tensor) shape for the network input. The TD and FD data are fed into the respective TD and FD networks, and a composite OOV Score is calculated from the TD and FD networks’ outputs, thus. Real and imaginary timepoints 10-to-1023 from the TD network’s OOV detection vector are added and divided by 1014; the initial 10 timepoints are ignored as OOV signals that originate at the top of the echo are indistinguishable from broad within-voxel MM and baseline signals^24^. Real and imaginary datapoints between 0.0 ppm and 4.1 ppm from the FD network detection vector are added and divided by the number of points. Finally, the TD and FD scores are averaged and multiplied by 100 to provide the composite OOV Score as a single integer that is written to a 4-byte binary file.

The 4-byte OOV Score is read from the binary results file by PEREGRINE (DAS-side), inside the VxWorks embedded RTOS and provided to the pulse sequence C++ code. OOV Scores are calculated every transient and sequence update decisions can occur immediately. For edited experiments, the sequence update decision is only made after a complete set of editing sub-experiments, based upon an overall OOV Score that is the average of the set of sub-experiment OOV Scores. When the OOV Score exceeds the user-defined threshold, the sequence is updated.

### 3.3. Real-Time Sequence Update

If an OOV signal is detected above the user-defined threshold, the crusher gradient scheme is updated to change the crushed k-space starting point of the OOV signal, cycling between 48 different geometries. These sample 6 pattern-to-axis mappings and independent positive-negative polarities for each gradient axis. For the pattern-axis mappings, the 3 different gradient patterns (Pattern-A, Pattern-B, and Pattern-C, as shown in Figure 2) are reordered (i.e., ABC, ACB, BAC, etc.) with respect to the sequence ‘MPS’ axes that correspond to the three slice-selective directions of the voxel. For the polarity, the gradient amplitudes can be positive or negative (e.g., +++, ++-, +-+, +--, etc.) on the M, P, and S axes.

**Figure 2.**
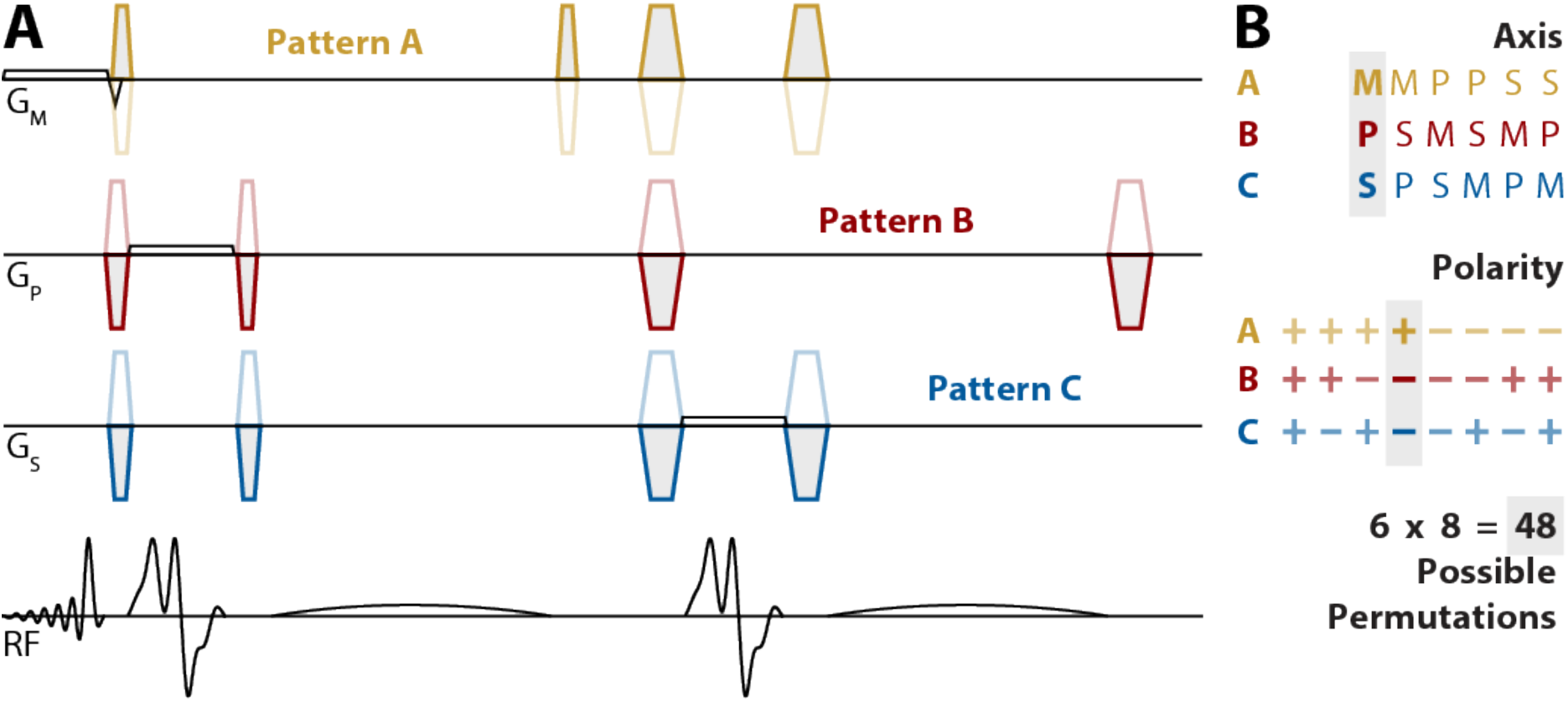
MEGA-PRESS Pulse Sequence with Corrective Gradients. **A)** MEGA-PRESS Pulse sequence implementation. The Crusher Gradient “Patterns” are shown in yellow for Pattern-A, red for Pattern-B, and blue for Pattern-C. These 3 patterns are used for each permutation, but can be assigned to different axes. The transparent gradients represent the ability of the for gradient’s amplitude to be positive or negative. **B)** The potential 6 axes and 8 polarities provide 48 different permutations.

If all 48 gradient options have been cycled through by OOV Scores exceeding threshold, the pulse sequence returns to a ‘recent-best’ gradient scheme. The pulse sequence tracks the lowest-score gradient scheme. The ‘recent-best’ index and value are updated, either: when a new lowest OOV Score is reached; or, if the ‘recent-best’ gradient scheme is the current gradient scheme, the OOV Score value is updated after each transient.

### 3.4. In Vivo Data

In vivo data were collected from 17 healthy subjects (33.67 ± 10.55 years; 12 females). All participants gave written informed consent before the scan session. A T1-weighted MPRAGE scan was acquired prior to MRS for voxel positioning (TR/TE = 8.1/3.7 ms; flip angle 8°; 1 mm in-plane resolution; slice thickness 1 mm; total scan time 2 min 47 s). A voxel of size 25 x 25 x 25 mm^3^ was localized to the medial prefrontal cortex (mPFC) as shown in Figure 3. Experiments were acquired on the Philips Achieva dStream scanner with dual-channel transmit body-coil and a 32-channel head receive coil. MRS data were acquired using a GABA-edited MEGA-PRESS (TE: 80 ms) pulse sequence with 20-ms editing pulses applied at 1.9 ppm and 7.46 ppm alternately^59–62^ and a VAPOR water suppression module of 722 ms duration^48^. These data were collected with 2048 time points, a spectral width of 2 kHz, and a TR of 2 s.

**Figure 3.**
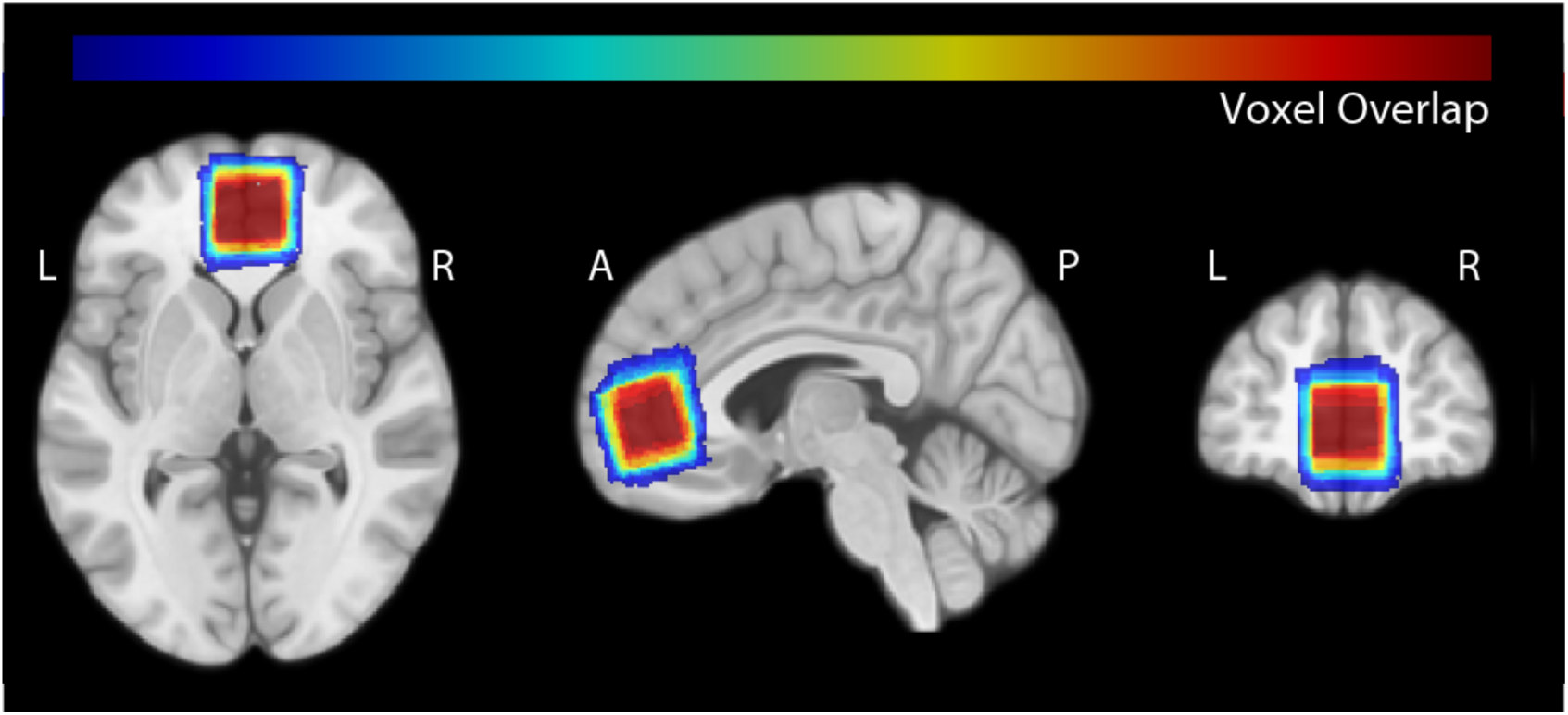
Voxel placement in the mPFC across participants. The native space voxel mask for each participant was normalized to MNI space and overlaid onto the spm152 template to display the anatomical overlap in voxel placement between participants. Warmer colors denote greater overlap.

Two spectroscopy experiments were collected, repeating full scan preparation between the two, with PEREGRINE running in order to provide OOV Scores for every transient. In the first experiment, the OOV Score was calculated without updating the sequence (referred to as AI-off), and in the second, the gradient scheme was updated, as described above (referred to as AI-on). Each experiment was collected with 192 transients (96 editing pairs). The OOV Score threshold to trigger sequence update was set to 15; this was chosen empirically based upon preliminary testing and prior work.

We assess: 1) whether real-time AI can successfully identify gradient schemes that reduce OOV artifacts; 2) whether dynamic sequence updates result in reduced OOV artifacts overall; and 3) whether overall quality of linear combination model fitting of spectra is improved by dynamic sequence updates. Because the sequence-update algorithm is not triggered for transients with sub-threshold OOV Scores, the algorithm tends to dwell on gradient schemes with better OOV characteristics. We assess (1) by defining the ‘Dwell’ condition as all gradient schemes for which the algorithm acquired three or more consecutive editing pairs. In order to maintain an appropriate comparison, the transient indices identified for the ‘Dwell’ condition of the AI-on experiment are also used to define the ‘Dwell’ comparator for the AI-off experiment in the same subject providing a matched subset of data from the unchanging gradient permutation. We assess (2) by defining the ‘Full’ condition as all transients within the scan (regardless of gradient permutation). We assess (3) with the Full condition using Osprey^63^ (version 2.9.6) to calculate the Fit Quality Number (FQN), or the variance of the linear combination model’s residual divided by the variance of the noise^64^. In order to assess gradient permutations overall, the mean OOV Score was calculated from the AI-on scan for each gradient permutation across all subjects.

A Shapiro-Wilk Test^65^ was used to determine if data were normally distributed. Paired t-Tests^66^ were used to compare the OOV Scores between the AI-on and AI-off scans (for the ‘Dwell’ and ‘Full’ conditions separately). A Wilcoxon Signed-Rank Test^67^, which is less influenced by outliers, was used to compare the non-normally distributed FQN scores between the AI-on and AI-off scans.

## 4. Results

### 4.1. Neural Network Performance

The TD network, developed in Torch, achieved a median Dice coefficient of 0.967 (0.872–0.983 interquartile range) osn the testing set, replicating the previous results seen with the TensorFlow implementation^24^. The FD network achieved a median Dice coefficient of 0.964 (0.889–0.983 interquartile range). Across the test set of 9,600, the combined networks achieved a median difference (predicted -ground truth) OOV Score of 1 (0–2 interquartile range).

### 4.2. PEREGRINE

Each acquired transient successfully moved through the reconstruction pipeline and was streamed using a TCP socket. Data were then received by PEREGRINE, processed, passed through the networks, and an OOV Score was written (host-side) and read (DAS-side) for each individual transient for the AI-on and AI-off scans for each of the 17 subjects (6,528 transients total). On the scanner CPU, each network within the PEREGRINE application executed in 10 ms, on average.

### 4.3. In Vivo Data

Examples of in vivo transients and the neural network OOV detection vectors are shown in Figure 4. The OOV Scores across the experiment for the pair of edit-ON and edit-OFF transients is plotted in Figure 5A. Note that the algorithm successfully updates the gradient scheme after an OOV Score that is above threshold, and not when the score is below threshold. The algorithm detected an OOV Score above threshold in 79% of editing pairs sampled during the cycling portion of the experiment. As all subjects, except one, completed cycling, the average number of gradient permutation updates was 47.94. On average, subjects completed the full cycling period after 63.31 ± 7.91 editing pairs. Amongst the Dwell periods while cycling, the median length for which the experiment remained unchanged was 3 edit-ON/edit-OFF pairs (3 – 5 inter-quartile range), or 6 consecutive transients. Over the entire experiment, the Dwell condition accounted for 50.86% of the total scan.

**Figure 4.**
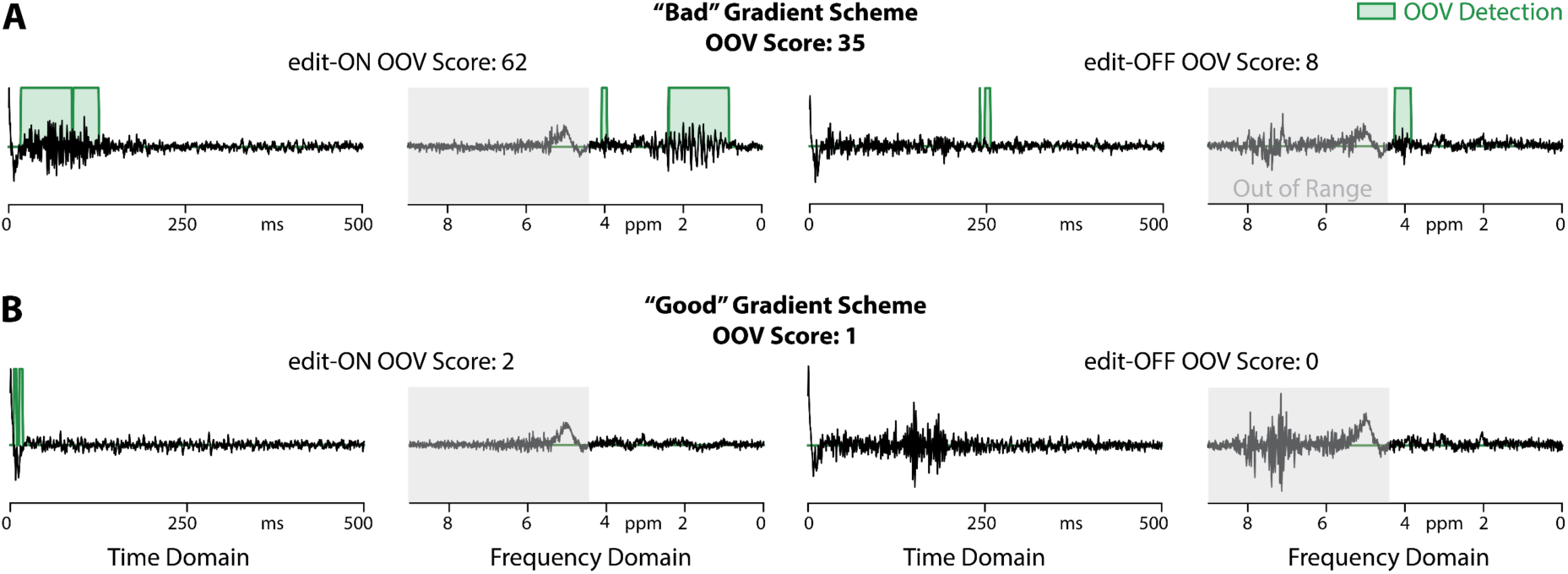
Examples of representative “Bad” and “Good” gradient permutations. **A)** A single transient example from a “Bad” gradient scheme permutation for both edit-ON and edit-OFF that produced an OOV Score above the user-defined threshold. **B)** A single transient example from a “Good” gradient permutation for both edit-ON and edit-OFF that produced an OOV Score under threshold. The input data is shown in black, neural network detection vector in green, and grey regions represent areas out of range (not included in OOV score calculation).

**Figure 5.**
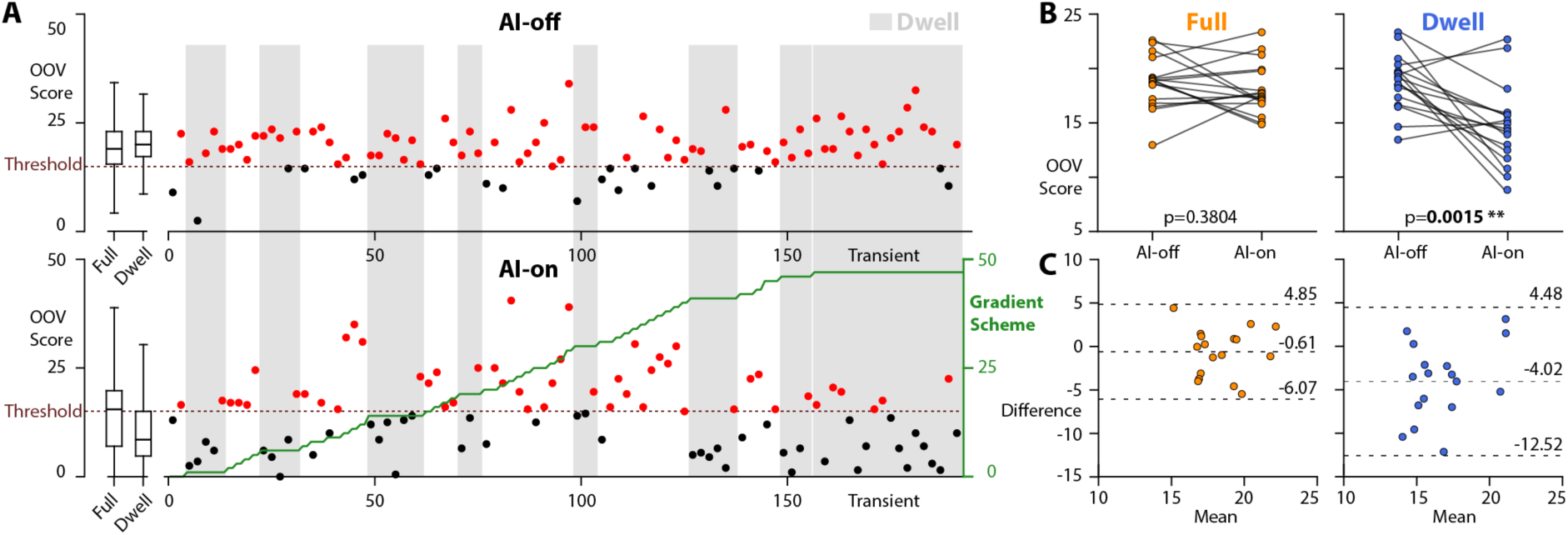
OOV Scores and Updates. **A)** The OOV Score for each MEGA-edited pair of transients across the AI-off (top) and AI-on (bottom) session. Red data points represent OOV scores above the threshold of 15 while black data points represent OOV Scores below threshold. The green line shows the gradient permutation across the AI-on session. **B)** AI-off compared to AI-on for the Full (orange) and the Dwell (blue) Conditions. **C)** Bland-Altman plots showing the difference in OOV scores for the Full (orange) and Dwell (blue) conditions. The center dashed lines represent the bias while the outer lines represent the upper and lower limits of agreement.

The mean OOV Scores for the Dwell condition were significantly lower for the AI-on experiment (14.70 ± 3.60) compared to the AI-off experiment (18.72 ± 2.49), with a t-statistic of 3.82 and p-value of 0.0015, as shown in Figure 5B. The OOV Scores for the Full condition had similar levels of OOV contamination between the AI-on (18.08 ± 2.29) and AI-off (18.69 ± 2.33) experiments, with a t-statistic of 0.90 and p-value of 0.38, as shown in Figure 5B. Bland-Altman plots for these comparisons are shown in Figure 5C.

Data modeling quality, as illustrated in Figure 6 and assessed by the FQN, was non-normally distributed according to the Shapiro-Wilk Test with a test statistic of 0.38 and a p-value of 1.4x10^-7^. The FQN was significantly lower for the AI-on experiment (2.31 ± 0.63) than for the AI-off experiment (4.49 ± 6.96), with a Wilcoxon Signed-Rank Test statistic of 31 and a p-value of 0.03 (as shown in Figure 7).

**Figure 6.**
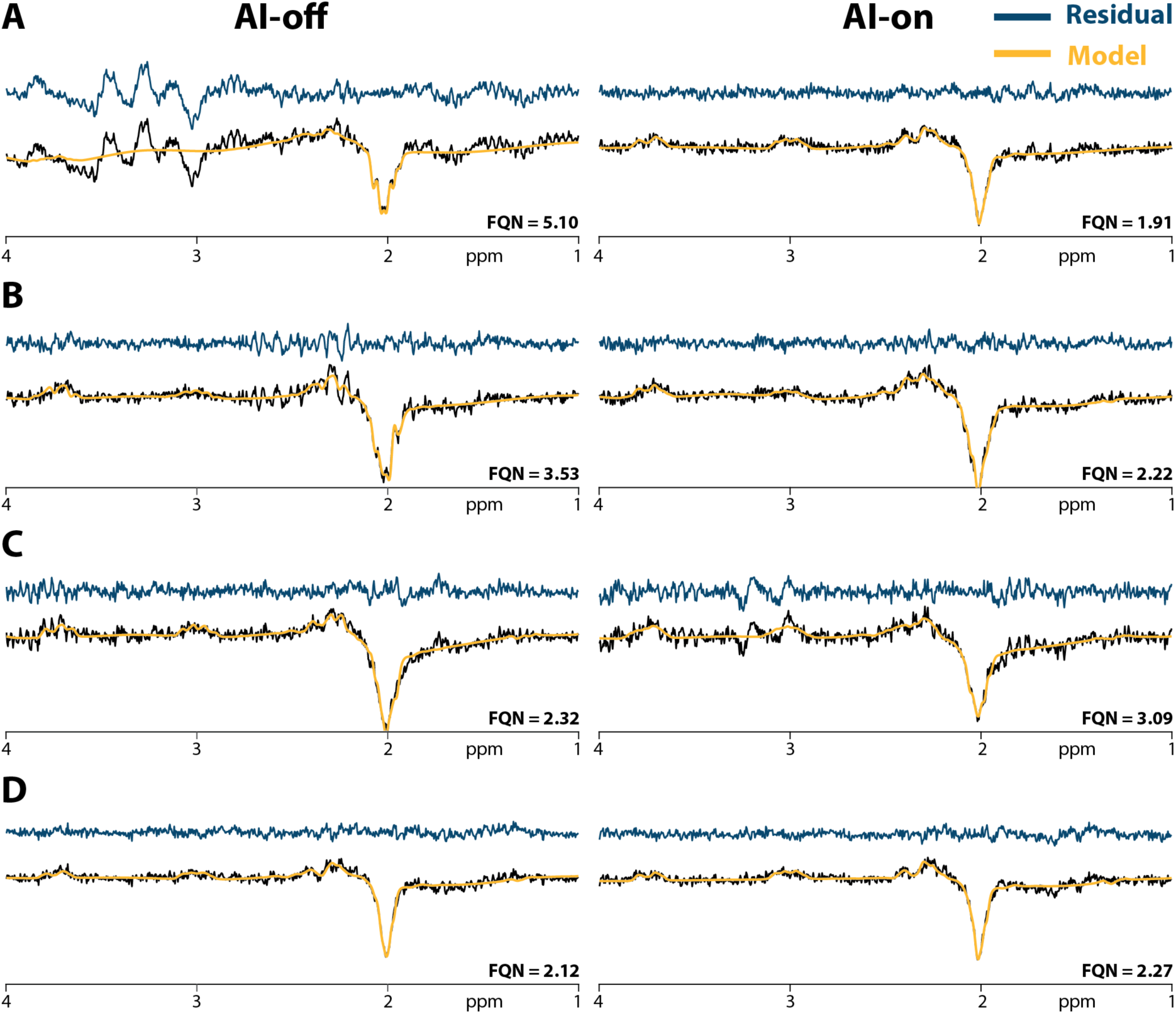
Linear Combination Model Fit for the AI-off and AI-on scans. **A)** and **B)** show poor model fit for the AI-off resulting from OOV signal and subtraction artifacts obstructing metabolite peaks. **C)** demonstrates a case where AI-on was markedly worse than AI-off due to cycling over gradient permutations that introduced worse artifacts than the default permutation experienced during the AI-off. **D)** demonstrates a case where both AI-off and AI-on were of good quality with relatively little OOV present.

**Figure 7.**
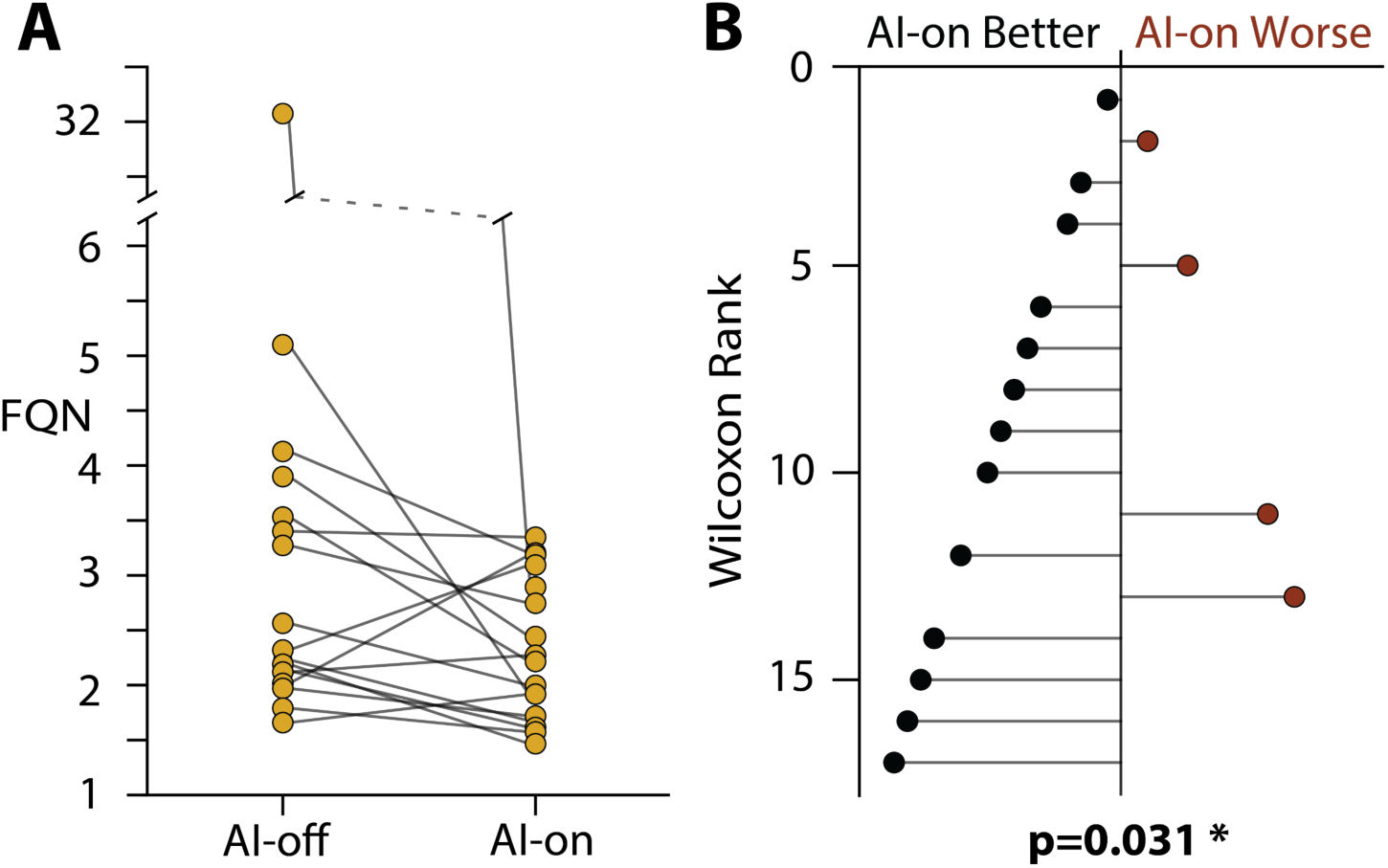
Fit Quality Number for the AI-off and AI-on scans. **A)** Comparison of the FQN for the AI-off and AI-on scans. **B)** The rankings from the Wilcoxon Signed-Rank Test are shown for the difference in FQN (AI-on – AI-off). This rank-based method is robust to outliers by testing position rather than the magnitude of the difference.

## 5. Discussion

This manuscript presents the first use of AI in real-time to assess acquired MR signals and adjust acquisition parameters of an ongoing scan accordingly. We introduced PEREGRINE for per-transient neural network assessment of spectral features with real-time sequence adjustment based upon the results, or put more simply, AI-based data-driven OOV detection and real-time gradient updates within an MRS experiment to minimize OOV artifacts in the MRS data. To achieve real-time AI-based detection of OOV signals, the entire workflow had to run between the end of one data acquisition window and the start of the upcoming TR. Importantly, the real-time components were fit within the constraints of typical MRS protocols, namely a TR of 2000 ms. This leaves roughly 200 ms between the end of one acquisition period and the beginning of the next water suppression module to successfully transfer data from the spectrometer to the reconstructor, run pre-processing, perform the forward pass through the neural networks, and make a decision based on the result. Without fail, PEREGRINE was able to evaluate the data, generate an OOV Score, and make sequence updates. This is an important proof-of-principle for within experiment AI assessment of data quality and corrective adjustments.

Overall, the AI-on scan produced improved spectra compared to the AI-off scan in terms of FQN. Interestingly, while searching for the best gradient permutation, the AI-on scan iterated over gradient schemes that were worse than the default (i.e., higher OOV Scores) and the mean OOV Score was comparable across both Full scans. The default gradient permutation 0 was not chosen randomly, but inherited from previous efforts to address OOV contamination, and ranked 10^th^ out of 48 within the full set of gradient permutations. This ‘better-than-average’ permutation for the AI-off control also mitigates against an improvement in average OOV Scores overall. The ability of PEREGRINE to select and stay on high-quality gradient permutations across the session, was supported by the lower mean OOV Score in the Dwell condition. Note that there is some potential selection bias when comparing OOV Scores for the Dwell condition, but that end result of this selection is improved data quality overall as judged by the FQN in the Full condition.

One negative impact of OOV signals on the processing workflow is in biasing shot-to-shot frequency-and-phase correction (FPC). Most FPC algorithms assume that the transients all contain the same information, and only differ by a phase or frequency shift. This assumption is violated in the presence of OOV signals that do not follow the intended coherence transfer pathway, because they have a different phase representation across the transients to the metabolite signals. This means that datasets containing OOV artifacts tend to be more misaligned than high-quality data, leading to greater subtraction artifacts. This secondary effect of OOV signals strongly contributes to the FQN results, in addition to the direct presence of OOV artifacts in the spectra. Indeed, in the two datasets that showed markedly worse FQN for AI-on than for AI-off, this loss of alignment drove the outcome.

At the outset, the expectation was that some gradient permutations would provide data that were free from OOV signals for each subject. Therefore, an algorithm was set up that would cycle rapidly through permutations until a desirable uncontaminated permutation was identified. It is a surprising finding, and an important one, that no single gradient permutation fully removed all OOV signals. Regardless of gradient scheme, OOV signals were detected in 79% editing pairs within the cycling period. Clearly, the number of latent OOV signals, crushed out into k-space but potentially refocused by local field gradients, was much larger than previously appreciated, and more diverse in terms of refocusing direction. Amongst all subjects, the likelihood of detecting OOV signals (and thus cycling to a new scheme) was high, and even below-threshold scores (where the algorithm dwelled) were often only just below threshold and followed by an above-threshold score. There *were* gradient permutations with low OOV contamination, but none that consistently returned an OOV Score below threshold. Thus, all but one subject’s AI-on experiment cycled through all 48 gradients permutations. Note that while geometric changes to the gradient scheme impact OOV signals, they are not expected to substantially influence metabolite signal localization, amplitude, or editing efficiency.

In designing an algorithm to search through the gradient permutations, we prioritized the ability to move on quickly from a ‘bad’ permutation, with a relatively low threshold. In this aspect, the algorithm was successful. With hindsight afforded by the presented data, selecting a different algorithm that encourages persistence with good permutations may have been ideal. For example, the ‘switch’ signal could be dependent upon the average OOV Score for the current permutation, which would make each subsequent transient less influential and cause the algorithm to dwell longer on relatively good options. An alternative algorithm would be to reject above-threshold transients and acquire additional transients to replace them. This would likely result in higher quality data overall at the expense of increased scan time, a delicate balance that would require careful consideration of the threshold.

One factor that was also poorly understood is the consistency of OOV signals across transients – in vivo, it is known that they are less stable than in-voxel metabolite signals. Many experimental factors, including motion, B0 drift, water suppression instability, etc., could also influence the OOV Score. Relying upon an average score threshold would make the experiment less rapidly responsive if the experiment switched state such that a previously good permutation became less favorable. Uncertainty over the consistency of signals also led us to default to the ‘recent best’ option upon exhausting all permutations, rather than the global best – a majority of scans settled on permutations 47 and 48, which were among the top 5 permutations, but not the best. Selecting the global best, even if it was sampled several minutes ago, represents a gamble on the consistency of the experiment that may or may not be warranted. Future work will investigate better algorithms, including algorithms that themselves apply AI to the decision.

Real-time AI is an exciting frontier of MRS that has the potential to address a longstanding weakness of the inconsistent data quality which often cannot easily be identified at the time of scanning. PEREGRINE was successfully deployed for the proof-of-principle task of OOV detection and gradient permutation, demonstrating the potential for alternative applications. The speed of AI and noise-tolerance for single transients make it possible to investigate aspects of the experiment that would previously have been impossible to fully sample. Encoding expert-level data assessment within the sequence addresses a further barrier – access to expertise. Every MR spectroscopist has experienced the disappointment of coming to analyze a research cohort, only to discover that some, or all, subjects’ data are of low quality. AI-embedded acquisition move this expert review process onto the scanner, while the subject is available, so that the experiment can be adjusted to acquire better-quality data.

## CRediT

**Aaron T. Gudmundson:** Conceptualization, Data Curation, Formal Analysis, Investigation, Methodology, Project administration, Software, Visualization, Writing-original draft, and Writing-Review & Editing. **Zahra Shams:** Conceptualization, Data Curation, Investigation, Methodology, Resources, Software, and Writing-Review & Editing. **Abdelrahman Gad:** Investigation, Resources, Writing-original draft, Writing-Review &Editing. **Shuyuan Wang:** Investigation and Writing-Review & Editing. **Dunja Simicic:** Investigation, Resources, Writing-Review &Editing. **Saipavitra Murali-Manohar:** Investigation, Resources, and Writing-Review &Editing. **Gizeaddis L. Simegn:** Formal Analysis, Visualization, Writing-Review & Editing. **Ipek Özdemir:** Software and Writing-Review & Editing. **Christopher W. Davies-Jenkins:** Formal Analysis and Writing-Review & Editing. **Yulu Song:** Investigation, Resources. **Vivek Yedavalli:** Investigation and Resources. **Georg Oeltzschner:** Conceptualization, Supervision, and Writing-Review & Editing. **Omer Burak Demirel:** Software and Supervision. **Jeremias Sulam:** Supervision, and Writing-Review & Editing. **Michael Schär:** Software, Supervision, and Writing-Review & Editing. **Sandeep Ganji:** Software, Supervision, and Writing-Review & Editing. **Richard A. E. Edden:** Conceptualization, Funding acquisition, Methodology, Resources, Supervision, Visualization, Writing-original draft, and Writing-Review & Editing.

## Acknowledgements

We acknowledge the support and contributions of Spencer Waddle, Melvyn Ooi, and Ryan Robison from Philips.

This work has been supported by the National Institute of Health, grants K99 HD118185, K99 EB034768, R21 EB033516, R01 EB035529, R01 EB016089, R01 EB023963, R01 EB032788, and P41 EB031771.

## Appendix

**Appendix 1:** MRSinMRS^68^.

| **1. Hardware** |  |
|---|---|
| a. Field strength [T] | 3 T |
| b. Manufacturer | Philips |
| c. Model (software version if available) | Achieva dStream (R57) |
| d. RF coils: nuclei (transmit/receive), number of channels, type, body part | <sup>1</sup>H, dual-channel body-coil transmit, 32-channel and 32-channel head coil receive |
| e. Additional hardware | N/A |
| **2. Acquisition** |  |
| a. Pulse sequence | MEGA-PRESS |
| b. Volume of Interest (VOI) locations | Prefrontal Cortex (PFC) |
| c. Nominal VOI size [cm<sup>3</sup>] | 2.5 x 2.5 x 2.5 cm<sup>3</sup> |
| d. TR/TE [ms] | TR 2000 ms / TE 80 ms |
| e. Total number of Excitations or acquisitions per spectrum | 192 transients |
| f. Additional sequence parameters (spectral width in Hz, number of spectral points) | 2000 Hz, 2048 points |
| g. Water Suppression Method | VAPOR |
| h. Shimming Method, reference peak, and thresholds for “acceptance of shim” chosen | B<sub>0</sub> shimming was performed in the selected voxels by second-order projection-based shimming (Philips pencil-beam). Reference peak: H<sub>2</sub>O |
| i. Triggering or motion correction method (respiratory, peripheral, cardiac triggering, incl. device used and delays) | N/A |
| **3. Data analysis methods and outputs** |  |
| a. Analysis software | Osprey (version 2.9.6) |
| b. Processing steps deviating from quoted reference or product | - Frequency-and-phase alignment of the individual transients via probabilistic spectral registration.<br>- Alignment of the averaged sub-spectra using Osprey’s “L2-norm” option.<br>- Residual water signal was removed using a Hankel singular value decomposition (HSVD) filter. |
| c. Output measure | Residual of modeling and residual of noise |
| d. Quantification references and assumptions, fitting model assumptions | Osprey Parameters:<br>- MM3coModel = “3to2MM” which determined the parametrization of the co-edited 3-ppm. macromolecular signal (here constrained to the 0.9-ppm MM signal).<br>- FWHMMM3Co = 14 the width of the co-edited MM signal in Hz.<br>- bLineKnotSpace = 0.4: cubic spline knot spacing. |
| **4. Data Quality** |  |
| a. Reported variables | SNR and linewidth not described |
| b. Data exclusion criteria | No subjects excluded |
| c. Quality measures of postprocessing Model fitting | Fit Quality Number (FQN) |
| d. Sample Spectrum | Figures 4 and 6 |

## Abbreviations

AGNOSTIC: Adaptable Generalized Neural-Network Open-source Spectroscopy Training dataset of Individual Components
AI: Artificial Intelligence
API: Application Programming Interface
DAS: Data Acquisition System
DOTCOPS: Dephasing Optimization Through Coherence Order Pathway Selection
FD: Frequency-domain
FPC: Frequency and Phase Correction
FQN: Fit Quality Number
mPFC: Medial Prefrontal Cortex
OOV: Out-of-voxel
PEREGRINE: Per Excitation Realtime Execution & Guided Responses with Integrated Neural-network Evaluation
PRIME: Philips Reconstruction Injector Modulator and Emitter
RTOS: Realtime Operating System
SOS: Standard Operating System
TCP: Transmission Control Protocol
TD: Time-domain

## References

1. Barker PB, Lin DDM. In vivo proton MR spectroscopy of the human brain. Prog Nucl Magn Reson Spectrosc. 2006;49(2):99–128. doi:10.1016/j.pnmrs.2006.06.002

2. Wilson M, Andronesi O, Barker PB, et al. Methodological consensus on clinical proton MRS of the brain: Review and recommendations. Magn Reson Med. 2019;82(2):527–550. doi:10.1002/mrm.27742

3. Keating B, Deng W, Roddey JC, et al. Prospective motion correction for single-voxel 1H MR spectroscopy. Magn Reson Med. 2010;64(3):672–679. doi:10.1002/mrm.22448

4. Marsman A, Lind A, Petersen ET, Andersen M, Boer VO. Prospective frequency and motion correction for edited 1H magnetic resonance spectroscopy. NeuroImage. 2021;233:117922. doi:10.1016/j.neuroimage.2021.117922

5. Andrews-Shigaki BC, Armstrong BSR, Zaitsev M, Ernst T. Prospective motion correction for magnetic resonance spectroscopy using single camera retro-grate reflector optical tracking. J Magn Reson Imaging. 2011;33(2):498–504. doi:10.1002/jmri.22467

6. Lange T, Maclaren J, Buechert M, Zaitsev M. Spectroscopic imaging with prospective motion correction and retrospective phase correction. Magn Reson Med. 2012;67(6):1506–1514. doi:10.1002/mrm.23136

7. Zaitsev M, Speck O, Hennig J, Büchert M. Single-voxel MRS with prospective motion correction and retrospective frequency correction. Published online 2010. doi:10.1002/nbm.1469

8. Adanyeguh IM, Bikkamane Jayadev N, Henry PG, Deelchand DK. Fast high-resolution prospective motion correction for single-voxel spectroscopy. Magn Reson Med. 2024;91(4):1301–1313. doi:10.1002/mrm.29950

9. Deelchand DK, Joers JM, Auerbach EJ, Henry PG. Prospective motion and B0 shim correction for MR spectroscopy in human brain at 7T. Magn Reson Med. 2019;82(6):1984–1992. doi:10.1002/mrm.27886

10. Hess AT, Dylan Tisdall M, Andronesi OC, Meintjes EM, van der Kouwe AJW. Real-time motion and B0 corrected single voxel spectroscopy using volumetric navigators. Magn Reson Med. 2011;66(2):314–323. doi:10.1002/mrm.22805

11. Hess AT, Andronesi OC, Dylan Tisdall M, Gregory Sorensen A, van der Kouwe AJW, Meintjes EM. Real-time motion and B0 correction for localized adiabatic selective refocusing (LASER) MRSI using echo planar imaging volumetric navigators. NMR Biomed. 2012;25(2):347–358. doi:10.1002/nbm.1756

12. Hess AT, Jacobson SW, Jacobson JL, Molteno CD, van der Kouwe AJW, Meintjes EM. A comparison of spectral quality in magnetic resonance spectroscopy data acquired with and without a novel EPI-navigated PRESS sequence in school-aged children with fetal alcohol spectrum disorders. Metab Brain Dis. 2014;29(2):323–332. doi:10.1007/s11011-014-9487-6

13. Keating B, Ernst T. Real-time dynamic frequency and shim correction for single-voxel magnetic resonance spectroscopy. Magn Reson Med. 2012;68(5):1339–1345. doi:10.1002/mrm.24129

14. Beracha I, Seginer A, Tal A. Adaptive model-based Magnetic Resonance. Magn Reson Med. 2023;90(3):839–851. doi:10.1002/mrm.29688

15. Bjørkeli EB, Geitung JT, Esmaeili M. Artificial intelligence-powered four-fold upscaling of human brain synthetic metabolite maps. J Int Med Res. 2025;53(4):03000605251330578. doi:10.1177/03000605251330578

16. Bugler H, Berto R, Souza R, Harris AD. Frequency and phase correction of GABA-edited magnetic resonance spectroscopy using complex-valued convolutional neural networks. Magn Reson Imaging. 2024;111:186–195. doi:10.1016/j.mri.2024.05.008

17. Chen D, Hu W, Liu H, et al. Magnetic Resonance Spectroscopy Deep Learning Denoising Using Few in Vivo Data. IEEE Trans Comput Imaging. 2023;9:448–458. doi:10.1109/TCI.2023.3267623

18. Dziadosz M, Rizzo R, Kyathanahally SP, Kreis R. Denoising single MR spectra by deep learning: Miracle or mirage? Magn Reson Med. Published online June 18, 2023. doi:10.1002/mrm.29762

19. Iqbal Z, Nguyen D, Hangel G, Motyka S, Bogner W, Jiang S. Super-Resolution 1H Magnetic Resonance Spectroscopic Imaging Utilizing Deep Learning. Front Oncol. 2019;9. doi:10.3389/fonc.2019.01010

20. Ma DJ, Le HAM, Ye Y, et al. MR spectroscopy frequency and phase correction using convolutional neural networks. Magn Reson Med. 2022;87(4):1700–1710. doi:10.1002/mrm.29103

21. Ma DJ, Yang Y, Harguindeguy N, et al. Magnetic Resonance Spectroscopy Spectral Registration Using Deep Learning. J Magn Reson Imaging. 2024;59(3):964–975. doi:10.1002/jmri.28868

22. Shamaei A, Starcukova J, Pavlova I, Starcuk Z. Model-informed unsupervised deep learning approaches to frequency and phase correction of MRS signals. Magn Reson Med. 2023;89(3):1221–1236. doi:10.1002/mrm.29498

23. Tapper S, Mikkelsen M, Dewey BE, et al. Frequency and phase correction of J-difference edited MR spectra using deep learning. Magn Reson Med. 2021;85(4):1755–1765. doi:10.1002/mrm.28525

24. Gudmundson AT, Davies-Jenkins CW, Özdemir İ, et al. Application of a 1H brain MRS benchmark dataset to deep learning for out-of-voxel artifacts. Imaging Neurosci. 2023;1:1-15. doi:10.1162/imag_a_00025

25. Gurbani SS, Schreibmann E, Maudsley AA, et al. A convolutional neural network to filter artifacts in spectroscopic MRI. Magn Reson Med. 2018;80(5):1765–1775. doi:10.1002/mrm.27166

26. Kyathanahally SP, Döring A, Kreis R. Deep learning approaches for detection and removal of ghosting artifacts in MR spectroscopy. Magn Reson Med. 2018;80(3):851–863. doi:10.1002/mrm.27096

27. Merkofer JP, van de Sande DMJ, Amirrajab S, Min Nam K, van Sloun RJG, Bhogal AA. WAND: Wavelet Analysis-Based Neural Decomposition of MRS Signals for Artifact Removal. NMR Biomed. 2025;38(6):e70038. doi:10.1002/nbm.70038

28. Lee HH, Kim H. Deep learning-based target metabolite isolation and big data-driven measurement uncertainty estimation in proton magnetic resonance spectroscopy of the brain. Magn Reson Med. 2020;84(4):1689–1706. doi:10.1002/mrm.28234

29. Rizzo R, Dziadosz M, Kyathanahally SP, Shamaei A, Kreis R. Quantification of MR spectra by deep learning in an idealized setting: Investigation of forms of input, network architectures, optimization by ensembles of networks, and training bias. Magn Reson Med. 2023;89(5):1707–1727. doi:10.1002/mrm.29561

30. Shamaei A, Starcukova J, Starcuk Z. Physics-informed deep learning approach to quantification of human brain metabolites from magnetic resonance spectroscopy data. Comput Biol Med. 2023;158:106837. doi:10.1016/j.compbiomed.2023.106837

31. LeCun Y, Bengio Y, Hinton G. Deep learning. Nature. 2015;521(7553):436-444. doi:10.1038/nature14539

32. Amos B. Tutorial on amortized optimization. arXiv. Preprint posted online October 5, 2025:arXiv:2202.00665. doi:10.48550/arXiv.2202.00665

33. Dvornik N, Shmelkov K, Mairal J, Schmid C. BlitzNet: A Real-Time Deep Network for Scene Understanding. In: 2017 IEEE International Conference on Computer Vision (ICCV). IEEE; 2017:4174-4182. doi:10.1109/ICCV.2017.447

34. Redmon J, Divvala S, Girshick R, Farhadi A. You Only Look Once: Unified, Real-Time Object Detection. In: 2016 IEEE Conference on Computer Vision and Pattern Recognition (CVPR). IEEE; 2016:779-788. doi:10.1109/CVPR.2016.91

35. Ernst T, Chang L. Elimination of artifacts in short echo time H MR spectroscopy of the frontal lobe. Magn Reson Med. 1996;36(3):462–468. doi:10.1002/mrm.1910360320

36. Kreis R. Issues of spectral quality in clinical1H-magnetic resonance spectroscopy and a gallery of artifacts. NMR Biomed. 2004;17(6):361–381. doi:10.1002/nbm.891

37. Moonen CTW, Sobering G, Van Zijl PCM, Gillen J, Von Kienlin M, Bizzi A. Proton spectroscopic imaging of human brain. J Magn Reson 1969. 1992;98(3):556-575. doi:10.1016/0022-2364(92)90007-T

38. Starck G, Carlsson A, Ljungberg M, Forssell-Aronsson E. k-space analysis of point-resolved spectroscopy (PRESS) with regard to spurious echoes in in vivo (1)H MRS. NMR Biomed. 2009;22(2):137–147. doi:10.1002/nbm.1289

39. Shams Z, Gad A, Gudmundson AT, et al. Identifying Out-of-Voxel Echoes in Edited MRS With Phase Cycle Inversion. Magn Reson Med. 2026;95(4):1921–1933. doi:10.1002/mrm.70211

40. Carlsson Å, Ljungberg M, Starck G, Forssell-Aronsson E. Degraded water suppression in small volume 1H MRS due to localised shimming. Magn Reson Mater Phys Biol Med. 2011;24(2):97–107. doi:10.1007/s10334-010-0239-2

41. Hurd RE. Gradient-enhanced spectroscopy. J Magn Reson 1969. 1990;87(2):422-428. doi:10.1016/0022-2364(90)90021-Z

42. Landheer K, Juchem C. Dephasing optimization through coherence order pathway selection (DOTCOPS) for improved crusher schemes in MR spectroscopy. Magn Reson Med. 2019;81(4):2209–2222. doi:10.1002/mrm.27587

43. Simegn GL, Shams Z, Murali-Manohar S, et al. Gradient Scheme Optimization for PRESS-Localized Edited MRS Using Weighted Pathway Suppression. NMR Biomed. 2026;39(1):e70182. doi:10.1002/nbm.70182

44. Song Y, Zöllner HJ, Hui SCN, Hupfeld KE, Oeltzschner G, Edden RAE. Impact of gradient scheme and non-linear shimming on out-of-voxel echo artifacts in edited MRS. NMR Biomed. 2023;36(2):e4839. doi:10.1002/nbm.4839

45. Bain AD. Coherence levels and coherence pathways in NMR. A simple way to design phase cycling procedures. J Magn Reson 1969. 1984;56(3):418-427. doi:10.1016/0022-2364(84)90305-6

46. Bodenhausen G, Kogler H, Ernst RR. Selection of coherence-transfer pathways in NMR pulse experiments. J Magn Reson 1969. 1984;58(3):370-388. doi:10.1016/0022-2364(84)90142-2

47. Haase A, Frahm J, Hanicke W, Matthaei D. 1H NMR chemical shift selective (CHESS) imaging. Phys Med Biol. 1985;30(4):341. doi:10.1088/0031-9155/30/4/008

48. Tkáč I, Starčuk Z, Choi IY, Gruetter R. In vivo 1H NMR spectroscopy of rat brain at 1 ms echo time. Magn Reson Med. 1999;41(4):649–656. doi:10.1002/(SICI)1522-2594(199904)41:4<649::AID-MRM2>3.0.CO;2-G

49. Chu A, Alger JR, Moore GJ, Posse S. Proton echo-planar spectroscopic imaging with highly effective outer volume suppression using combined presaturation and spatially selective echo dephasing. Magn Reson Med. 2003;49(5):817–821. doi:10.1002/mrm.10449

50. Duijn JH, Matson GB, Maudsley AA, Weiner MW. 3D phase encoding 1H spectroscopic imaging of human brain. Magn Reson Imaging. 1992;10(2):315–319. doi:10.1016/0730-725X(92)90490-Q

51. Shams Z, Klomp DWJ, Boer VO, Wijnen JP, Wiegers EC. Identifying the source of spurious signals caused by B0 inhomogeneities in single-voxel 1H MRS. Magn Reson Med. 2022;88(1):71–82. doi:10.1002/mrm.29222

52. Gudmundson AT, Koo A, Virovka A, et al. Meta-analysis and open-source database for in vivo brain Magnetic Resonance spectroscopy in health and disease. Anal Biochem. 2023;676:115227. doi:10.1016/j.ab.2023.115227

53. Paszke A, Gross S, Massa F, et al. PyTorch: An Imperative Style, High-Performance Deep Learning Library. *arXiv*. Preprint posted online December 3, 2019:arXiv:1912.01703. doi:10.48550/arXiv.1912.01703

54. Carass A, Roy S, Gherman A, et al. Evaluating White Matter Lesion Segmentations with Refined Sørensen-Dice Analysis. Sci Rep. 2020;10(1):1–19. doi:10.1038/s41598-020-64803-w

55. Dice LR. Measures of the Amount of Ecologic Association Between Species Author ( s ): Lee R . Dice Published by : Ecological Society of America Stable URL : http://www.jstor.org/stable/1932409. *Ecology*. 1945;26(3):297-302.

56. Sørensen T. A method of establishing groups of equal amplitude in plant sociology based on similarity of species and its application to analyses of the vegetation on Danish commons. K Dan Vidensk Selsk. 1948;5(4):1–34.

57. Demirel OB, Spencer Waddle, Ganji S, Ooi M, Robison R. An Open Innovation Emitter– Modulator–Injector Framework for Inline MRI Reconstruction (PRIME). In: 2026 ISMRM & ISMRT Annual Meeting & Exhibition. 2026.

58. Cooley JW, Tukey JW. An Algorithm for the Machine Calculation of Complex Fourier Series. Math Comput. 1965;19(90):297–301. doi:10.2307/2003354

59. Saleh MG, Rimbault D, Mikkelsen M, et al. Multi-vendor standardized sequence for edited magnetic resonance spectroscopy. NeuroImage. 2019;189:425–431. doi:10.1016/j.neuroimage.2019.01.056

60. Mescher M, Merkle H, Kirsch J, Garwood M, Gruetter R. Simultaneous in vivo spectral editing and water suppression. NMR Biomed. 1998;11(6):266–272. doi:10.1002/(SICI)1099-1492(199810)11:6<266::AID-NBM530>3.0.CO;2-J

61. Choi C, Ganji S, Hulsey K, et al. A comparative study of short-and long-TE ^1^H MRS at 3 T for *in vivo* detection of 2-hydroxyglutarate in brain tumors. doi:10.1002/nbm.2943

62. Murdoch JB, Lent AH, Kritzer MR. Computer-optimized narrowband pulses for multislice imaging. J Magn Reson 1969. 1987;74(2):226-263. doi:10.1016/0022-2364(87)90336-2

63. Oeltzschner G, Zöllner HJ, Hui SCN, et al. Osprey: Open-source processing, reconstruction C estimation of magnetic resonance spectroscopy data. J Neurosci Methods. 2020;343(June):108827. doi:10.1016/j.jneumeth.2020.108827

64. Kreis R, Boer V, Choi I, et al. Terminology and concepts for the characterization of in vivo MR spectroscopy methods and MR spectra: Background and experts’ consensus recommendations. NMR Biomed. 2021;34(5):e4347. doi:10.1002/nbm.4347

65. Shapiro SS, Wilk MB. An Analysis of Variance Test for Normality (Complete Samples). Biometrika. 1965;52(3/4):591–611. doi:10.2307/2333709

66. Rosner B. A Generalization of the Paired t-Test. J R Stat Soc Ser C Appl Stat. 1982;31(1):9–13. doi:10.2307/2347069

67. Woolson RF. Wilcoxon Signed-Rank Test. Accessed May 3, 2026. https://onlinelibrary.wiley.com/doi/10.1002/9780471462422.eoct979

68. Lin A, Andronesi O, Bogner W, et al. Minimum Reporting Standards for in vivo Magnetic Resonance Spectroscopy (MRSinMRS): Experts’ consensus recommendations. NMR Biomed. 2021;34(5):e4484. doi:10.1002/nbm.4484

